# Comparison of CcrM-dependent methylation in *Caulobacter crescentus* and *Brucella abortus* by nanopore sequencing

**DOI:** 10.1101/2024.03.01.583015

**Authors:** Maxwell Campbell, Ian Scott Barton, R. Martin Roop, Peter Chien

## Abstract

Bacteria rely on DNA methylation for restriction-modification systems and epigenetic control of gene expression. Here, we use direct detection of methylated bases by nanopore sequencing to monitor global DNA methylation in Alphaproteobacteria, where use of this technique has not yet been reported. One representative of this order, *Caulobacter crescentus*, relies on DNA methylation to control cell cycle progression, but it is unclear whether other members of this order, such as *Brucella abortus*, depend on the same systems. We addressed these questions by first measuring CcrM-dependent DNA methylation in *Caulobacter* and show excellent correlation between nanopore-based detection and previously published results. We then directly measure the impact of Lon-mediated CcrM degradation on the epigenome, verifying that loss of Lon results in pervasive methylation. We also show that the AlkB demethylase has no global impact on DNA methylation during normal growth. Next, we report on the global DNA methylation in *Brucella abortus* for the first time and find that CcrM-dependent methylation is reliant on Lon but impacts the two chromosomes differently. Finally, we explore the impact of the MucR transcription factor, known to compete with CcrM methylation, on the *Brucella* methylome and share the results with a publicly available visualization package. Our work demonstrates the utility of nanopore-based sequencing for epigenome measurements in Alphaproteobacteria and reveals new features of CcrM-dependent methylation in a zoonotic pathogen.

**Importance:** DNA methylation plays an important role in bacteria to maintain genome integrity and regulate of gene expression. We used nanopore sequencing to directly measure methylated bases in *Caulobacter crescentus* and *Brucella abortus*. In *Caulobacter*, we showed that stabilization of the CcrM methyltransferase upon loss of the Lon protease results in prolific methylation and discovered that the putative methylase AlkB is unlikely to have a global physiological effect. We measured genome-wide methylation in Brucella for the first time, revealing a similar role for CcrM in cell-cycle methylation but a more complex regulation by the Lon protease than in Caulobacter. Finally, we show how the virulence factor MucR impacts DNA methylation patterns in *Brucella*.

## Introduction

DNA methylation is a critical epigenetic modification where motif-specific addition of a methyl group to adenine or cytosine can affect gene regulation, structure, or properties (^1^). In bacteria, the methyltransferases responsible for this process are grouped into two main types: restriction-modification systems and orphan methyltransferases. While restriction-modification systems harbor DNA methyltransferases along with associated restriction enzymes that serve to protect the bacteria’s own DNA while degrading foreign DNA (^2,3^), orphan methyltransferases lack associated restriction endonucleases. Epigenetic marks laid down by orphan enzymes play crucial roles in controlling transcription in cellular processes such as initiation of DNA replication and development (^4,5^).

In the dimorphic alphaproteobacterium *Caulobacter crescentus (Caulobacter)*, the *ccrM* gene encodes an orphan cell cycle-regulated methyltransferase. CcrM generates N6-methyladenine (m6A) marks in GANTC motifs enriched in intergenic regions that often lead to changes in transcription of downstream genes. For instance, methylated GANTC sites in the *dnaA* promoter (^6^) or hemi-methylated GANTC sites in the *ctrA* promoter (^7^) increase the expression of these genes. Notably, CcrM is cell cycle regulated, with activity peaking at end of the S-phase of the cell cycle and then lost through rapid degradation by the Lon protease (^6,8^). Thus, newly synthesized DNA remains hemi-methylated during replication and full methylation occurs once the chromosome has fully replicated as CcrM activity peaks, resulting in a steady increase in GANTC methylation measuring from origin to terminus in asynchronous populations (^9,10^). Upregulation of CcrM activity due to overexpression (^10^) or stabilization (^8^) results in loss of the characteristic cell cycle GANTC methylation and an overall increase in methylation at CcrM-dependent sites. More recently, additional regulation of methylation has been proposed from findings that regions of hypomethylation seem to arise from loss of CcrM access to GANTC sites due to competitive binding of factors such as MucR (^11^).

Several outstanding questions emerge from these observations. For example, regulated removal of methylation marks by demethylases alters epigenetic status in eukaryotic genomes (^12^); however, demethylation of CcrM deposited methylation marks has not been shown. One particularly intriguing candidate for a possible demethylase is AlkB, a hydroxylase responsible for repair of alkylated DNA during damaging conditions (^13^). In *Caulobacter*, *alkB* expression is under cell cycle control, suggesting a possible role in this process (^14^). *In vitro*, *E. coli* AlkB is capable of removing m6A marks from DNA templates (^15^), leading to a tempting speculation that AlkB may contribute to *Caulobacter* genome methylation, perhaps during the cell cycle. In addition, although CcrM and Lon are highly conserved among Alphaproteobacteria, whether the model that Lon degradation of CcrM controls its activity as seen in *Caulobacter* extends to all Alphaproteobacteria remains unclear. One important example is in the zoonotic pathogen *Brucella abortus (Brucella)*, where both CcrM and Lon were found to be modulators of physiology and virulence (^16,17^), but to our knowledge no studies on methylation in this bacteria have been reported.

In this paper, we address some of these questions using nanopore-based sequencing to measure global methylation. We first establish that machine-based learning models and nanopore reads can accurately measure cell cycle dependent methylation in *Caulobacter*. We show that loss of Lon results in pervasive elevated methylation across all sites, as would be expected from the known elevated levels of CcrM in Δ*lon* strains and correlate these effects with known transcriptional changes. We test if AlkB controls methylation and while we find no global change, a handful of sites exhibit minor methylation marks that in at least one case alters gene expression level. Next, we profiled the global methylome of *Brucella* and found that both chromosomes show signatures of cell cycle and CcrM-dependent methylation. Interestingly, loss of Lon results in chromosome-specific loss of cell cycle methylation and global methylation of both chromosomes is lower, in contrast to the *Caulobacter* case. Finally, we address the impact of the MucR transcription factor on global methylation in *Brucella*.

## RESULTS

In *Caulobacter*, cell cycle regulated CcrM-dependent methylation controls DNA replication and gene expression (^18,19^). The prevailing model (Figure 1B) is that replication of the fully methylated chromosome initiates during the swarmer-stalk transition with newly synthesized DNA strands remaining unmethylated throughout replication (^9,10^). This process results in the genome transitioning from a fully methylated state to a hemi-methylated state as the replication fork progresses. Once replication is complete, CcrM activity surges due to increased expression and stabilization from Lon degradation resulting in full methylation of the chromosome (^9^). As a consequence, GANTC sites located nearer to the origin of replication are likely to remain hemi-methylated for a more extended period than those closer to the terminus.

**Figure 1:**
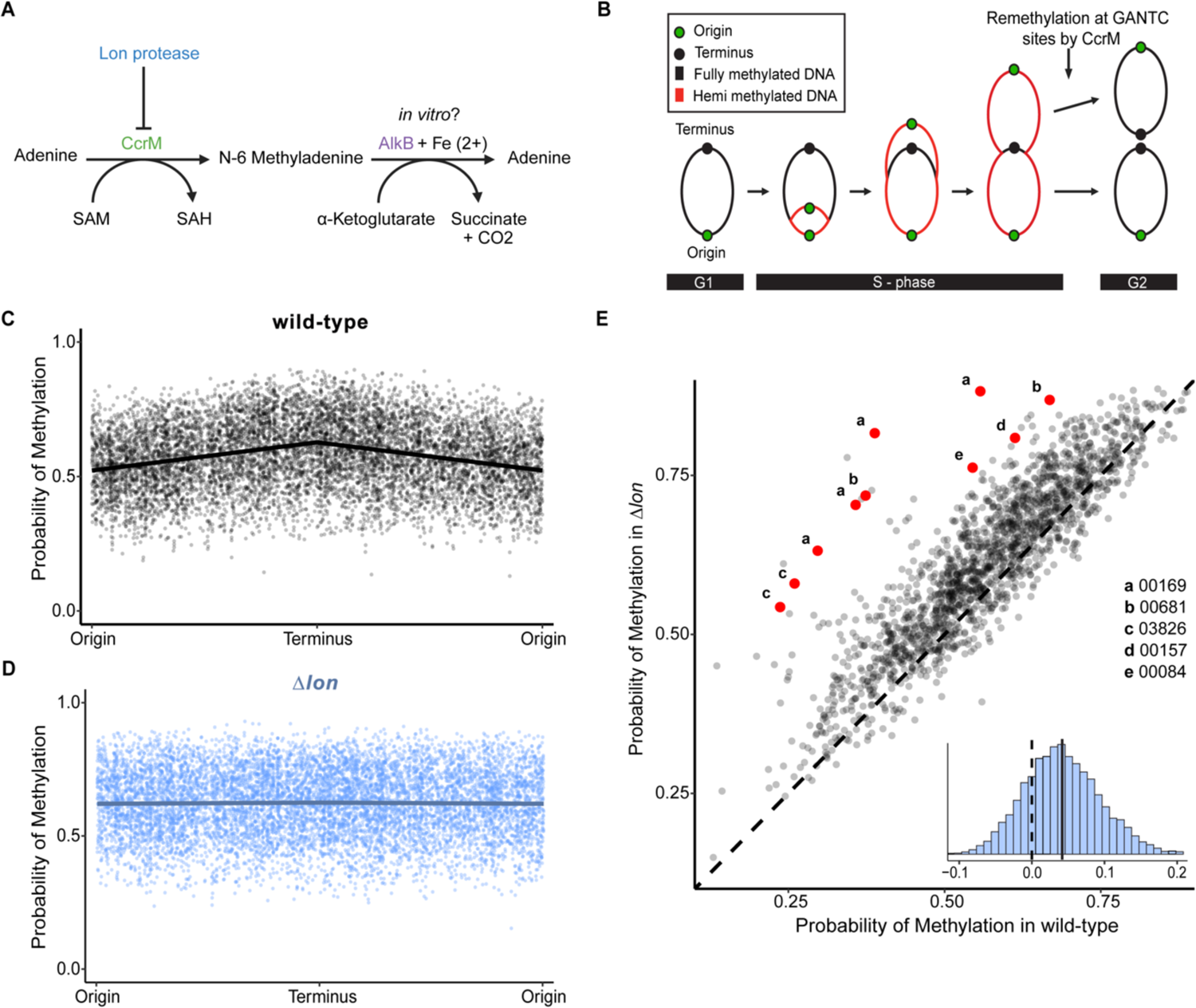
(**A**) CcrM facilitates the transfer of a methyl group from S-Adenosyl methionine (SAM) to adenine bases resulting in the formation of N-6 methyladenine and S-adenosyl-L-homocysteine (SAH) (^47^). Lon protease degrades CcrM, thereby restricting its activity to the predivisional cell state. The dealkylase AlkB could play a role in the removal of the m6A epigenetic mark as described *in vitro* (^15^). (**B**) The current model of chromosomal m6A CcrM-dependent methylation during replication. Newly synthesized DNA is unmethylated, resulting in a progressive transition into a fully to hemi-methylated state comparing origin to terminus until the expression of CcrM near the end of S-Phase. (**C, D**) Probability of CcrM-dependent methylation at GANTC motifs across the chromosome of *Caulobacter* in both wild-type (**C**) and Δ*lon* (**D**) strains. Lines are linear fits to methylation probability assuming symmetry over the terminus. **(E**) Scatter plot of probability of methylation at intergenic sites in Δ*lon* and wild type strains, with red dots marking sites with both significant transcriptional changes and deviating from the mean by 2.5σ (CCNA designation in legend; highlighted motifs listed in Table S4). Histogram inset, displays the distribution of differences between methylation of *Δlon* and wild-type with the dashed line indicating the line of identity and bold line displaying mean difference in methylation (Δ = 0.043).

Past global methylation studies of *Caulobacter* have used PacBio-based SMRT-Seq (^9,10^) and methylation-specific restriction enzymes (REC-Seq^11^). In this current work, we take advantage of nanopore sequencing combined with machine learning pipelines developed to call modifications of bases, including methylation. We first sequenced and called the methylome for wild-type *Caulobacter* using ONT v9.5 pores and mCaller (^20^). Our findings corroborated the expected methylation pattern, as GANTC methylation is low at the origin and gradually increases to maximum at the terminus, consistent with cell cycle dependent methylation and an asynchronous population of cells (Figure 1C). Our results align well with prior studies that employed SMRT sequencing (^10^), with strong correlations comparing average methylation across sections of the chromosome and somewhat weaker correlation between individual sites (SFig 1). Importantly, m6A marks that are not GANTC motifs, and therefore not subject to CcrM control, exhibit an even and consistently high methylation across the genome, supporting that CcrM is the only cell cycle regulated adenine methyltransferase (SFig 4). We conclude that methylome detection using nanopore sequencing can be used effectively for *Caulobacter* genomes.

### Loss of Lon globally affects CcrM-dependent methylation in *Caulobacter*

CcrM was the first substrate known to be degraded by the Lon protease in *Caulobacter* (^8^) and loss of Lon results in increased CcrM levels with increased methylation observed at individual sites (^8,21^). However, there has been no global measurement of methylation in Δ*lon Caulobacter*. We now address this gap by using nanopore sequencing. Consistent with a critical role of Lon degradation in controlling CcrM levels and timing, we find that the cell cycle-dependent variation in GANTC methylation is lost in Δ*lon* and methylation across the entire genome is elevated (Fig 1D). Again, this was limited to m6A methylation of GANTC as non-GANTC m6A marks remain the same as wild-type (SFigure S4). Based on this, Lon-dependent degradation of CcrM clearly plays a significant role in controlling CcrM activity globally.

Methylation of promoter regions is thought to affect transcription of the neighboring genes (^10,11,19,22^). We tested if this increase in methylation of promoter regions in Δ*lon* is correlated with changes in transcription in this strain. Using a cutoff of 300 base-pairs upstream of annotated genes as the promoter region, we performed a Fisher exact test comparing significant changes in gene expression between wild-type and Δ*lon* (^23^) to changes in methylation of those promoter regions. This reveals no significant enrichment (p= 0.206).

However, among the sites exhibiting methylation probabilities that diverged by more than 2.5 standard deviations from the adjusted mean in the Δ*lon* strain, five genes showed significant transcriptional alterations (CCNA_00169, CCNA_00681, CCNA_03826, CCNA_00157, and CCNA_00084; labeled in order as a-e in the figure) with differing numbers of CcrM-dependent methylation sites in the promoter regions (Figure 1E). We conclude that although there is not a global change in transcription directly attributable to changes in promoter methylation, some genes may be more susceptible than others.

### Loss of AlkB does not globally affect methylation

There is no *in vivo* evidence of demethylase activity to counter m6A modifications in bacteria. However, given that AlkB homologs have been shown to be biochemically proficient for removal of these modifications *in vitro* (^15^) we addressed whether AlkB affected CcrM-dependent methylation in *Caulobacter* using the same nanopore approach as before. There was a high correlation between methylation of Δ*alkB* and wild-type strains at all GANTC sites (Pearson coefficient 0.95). Differences in GANTC methylation across all the sites exhibited a normal distribution centered at zero (Anderson-Darling test), suggesting no significant global variation of methylation. We also tested whether AlkB might be affecting ‘spurious’ methylation of non-GANTC m6A sites, but found no significant difference in those comparisons either (SFigure 4).

Interestingly, one GANTC site that shows a higher degree of methylation in Δ*alkB* than expected (>2.5σ) is in the *ccrM* promoter region. This motif has been shown in past studies to have a minor influence on expression of *ccrM* (^24^). Based on western blots, we found that CcrM levels are approximately two-fold higher in Δ*alkB* strains compared to wild-type in early exponential phase (Figure 2) but reach the same final levels as cells enter stationary phase. We conclude that AlkB does not play a global role in genome methylation under standard lab conditions but may have a gene-specific impact as shown with *ccrM*.

**Figure 2:**
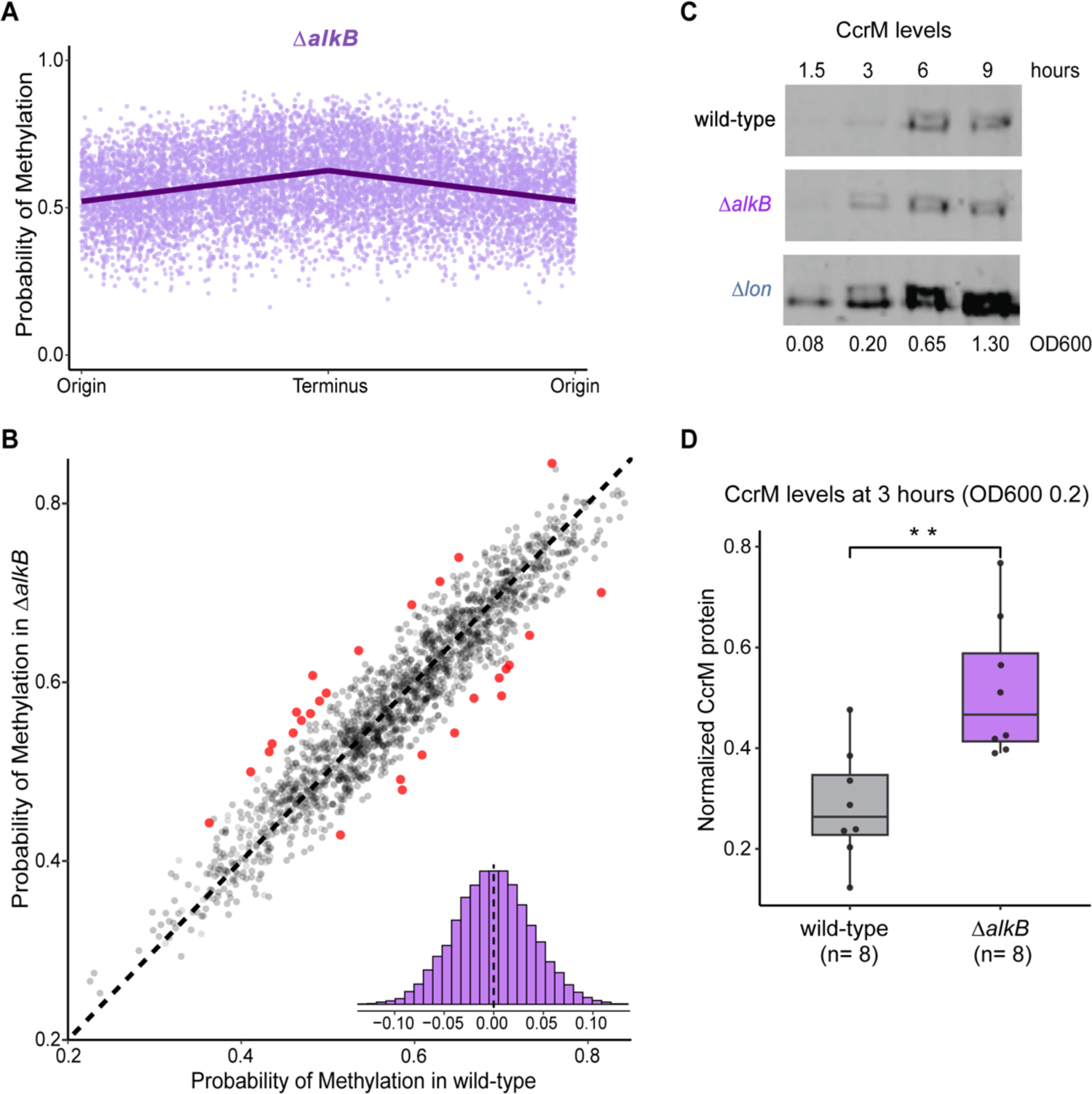
(A) Probability of methylation at GANTC motifs across the chromosome of *ΔalkB Caulobacter*. (B) Scatter plot comparing difference in probability of methylation at intergenic motifs between *ΔalkB* and wild-type *Caulobacter*, with points highlighted in red marking motifs that deviate from the mean by 2.5 standard deviations (List of highlighted motifs are available in Table S5). Histogram inset, displays the distribution of differences between all Δ*alkB* and wild-type motifs with the dashed line indicating the line of identity and mean. (C) Western blot of CcrM protein levels in wild-type (black/top), *ΔalkB* (purple/ middle), and *Δlon* (blue/bottom) during growth. (D) Quantification of CcrM protein levels after 3 hours of growth (OD600 0.2), in wild-type (black) and *ΔalkB* (purple) normalized to ClpP protein levels. (**) P-value of 0.002 by Student’s t-test.

### CcrM-dependent methylation in *Brucella* is cell cycle regulated

We next used a nanopore platform to determine the methylation status of the *Brucella* genome. In this case, newly released technology and pipelines from Oxford Nanopore Technology were used (^25^) for methylation detection due to changes by the sequencing vendor. We used wild-type, CcrM overexpressing (^17^), Δ*lon* (^16^), and Δ*mucR* (^18^) strains of *Brucella* to explore the molecular drivers of the methylome for this zoonotic pathogen.

Methylome measurements of wild-type *Brucella* reveal an increase in m6A at GANTC sites progressing from origin-proximal to terminal regions for both chromosomes (Fig 3 A), suggesting cell cycle dependent methylation. We reasoned that this was due to CcrM activity similar to *Caulobacter* as only m6A marks associated with GANTC motifs show this progressive change (SFig 5) and overexpression of CcrM results in increased methylation across the genome only at these cell cycle regulated GANTC sites (Fig 3A, B). Both chromosomes in *Brucella* showed the same degree of change in GANTC methylation suggesting that even though Chromosome II initiates replication after Chromosome I (^26^) both chromosomes are subject to the same cell cycle regulation by CcrM under normal conditions.

**Figure 3:**
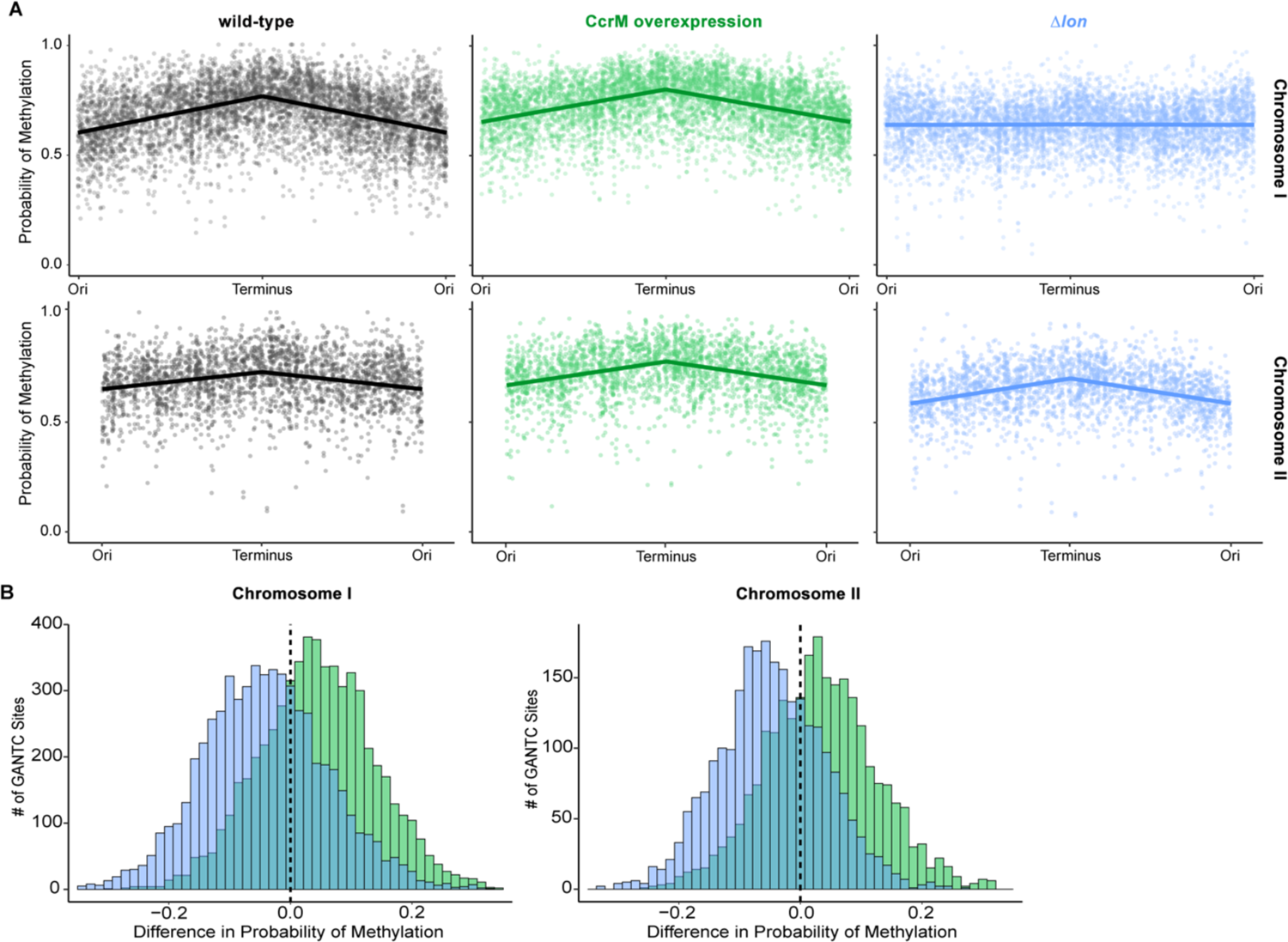
(**A**) Probability of CcrM-dependent methylation at GANTC motifs on Chromosome I (upper graphs) and Chromosome II (lower graphs) of *Brucella* in different genetic contexts: wild-type (black), with CcrM overexpression (green), and deletion of *lon* (blue). (**B**) Distribution of differences in strains where CcrM is overexpressed (green) or *Δlon* (blue) verse wild-type, across all GANTC sites and among both chromosomes (left: Chromosome I, right: Chromosome II)

**Figure 4:**
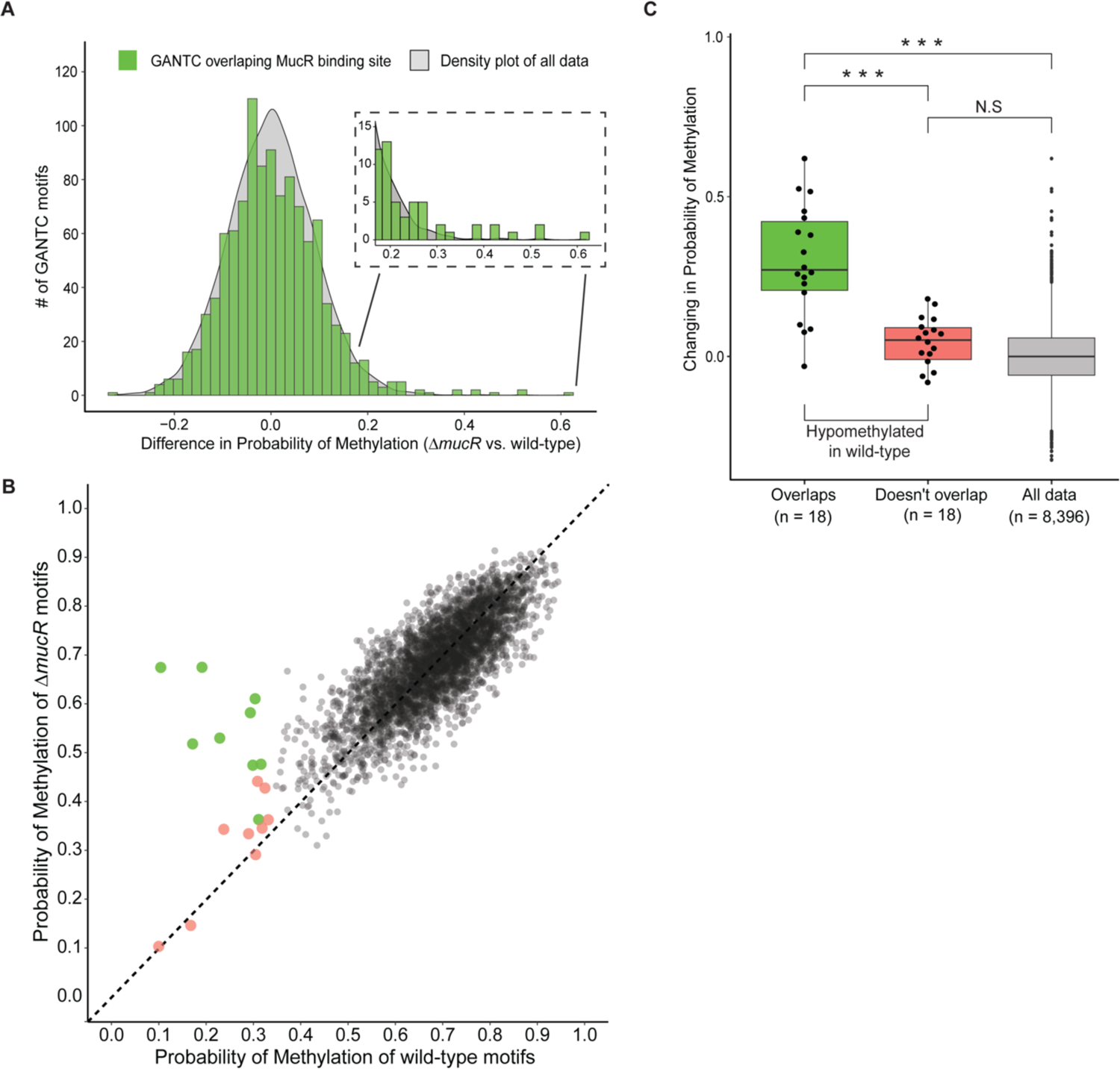
(**A**) Density plot (gray) depicting the distribution of differences in methylation probability between *ΔmucR* and wild-type strains at all GANTC motifs. Green histogram shows sites that overlap with MucR binding sites, with inset showing highest changing sites. (**B**) Scatterplot of methylation probability at GANTC sites across both chromosomes of *ΔmucR* and wild-type *Brucella*. Hypomethylated sites in the wild-type overlapping with MucR binding sites are highlighted in green, while those not overlapping are in red. (**C**) Box plots illustrating changes in methylation probability between *ΔmucR* and wild-type for sites that are hypomethylated in wild-type with (green) and without (red) overlap with MucR binding regions, along with all sites (gray). Student t-test shows p-value as 6.0 e-6 (***), 1.7e-10(***) and no significant difference (N.S).

We next explored the role of the Lon protease in CcrM dependent methylation in *Brucella*. Given the high conservation of both proteins (65-70% identity), we envisioned similar regulation. Surprisingly, we found substantial Lon-dependent methylation differences between these species. Specifically, Chromosome I showed the expected loss of cell cycle dependent methylation when *lon* was deleted, but Chromosome II did not (Figure 3A). In addition, upon loss of Lon both chromosomes exhibited lower methylation overall compared to the wild-type strain (Figure 3B). This contrasts the *Caulobacter* case where elevated CcrM-dependent methylation is seen when *lon* is deleted (Fig 1). The interpretation of these results is explored in the discussion, but it is clear that the link between Lon and CcrM is more complex in *Brucella*.

### MucR modulates local methylation in *Brucella*

In *Caulobacter*, the MucR transcription factor regulates developmental transitions (^27^) and occludes GANTC motifs methylated by CcrM (^11^). MucR also has important roles for virulence in *Brucella* (^28^) and recent ChIP-Seq/Hi-C experiments reveal that MucR may also play a role in structural organization of chromosomes (^18^). We investigated the role of MucR in methylation of *Brucella* chromosomes by measuring the methylome of a Δ*mucR* strains. Overall, there was no global difference in methylation between wild-type and Δ*mucR* genomes across either chromosome (Figure 4A). However, when we specifically focused on MucR binding sites (^18^) we found that some overlapping sites had unusually high methylation in the Δ*mucR* strain compared to wild-type (Figure 4A). Interestingly these sites are part of a cluster of 18 hypomethylated sites in wild-type genomes (Figure 4B). Of those hypomethylated sites, those that overlap with MucR binding sites show high methylation in Δ*mucR* while those that do not overlap with MucR binding sites are not affected (Figure 4C; Table S6). We conclude that MucR shields a subset of hypomethylated sites from methylation by CcrM, similar to the *Caulobacter* system, but does not universally protect binding sites from methylation.

## Discussion

Epigenetic modification of bacterial chromosomes was once considered only for restriction-modification systems used to protect cells from phage attack. The identification of orphan methyltransferases that modify chromosomes during development suggested that these modifications could also serve regulatory roles, much like CpG methylation in eukaryotes. Bisulfite-sequencing is a common method for detecting cytosine methylation in eukaryotic genomes; however, the primary regulatory modification in bacteria is adenine methylation, which cannot be converted in the same manner as methyl-cytosine. Use of single-molecule techniques such as Pacific Biosciences SMRT sequencing platform has resulted in direct observation of DNA methylation and application of this to bacterial genomes has made inroads into this regulatory mechanism. However, the costs associated with this approach make it challenging to do routinely. In this work, we take advantage of machine learning pipelines and increasingly inexpensive nanopore sequencing to measure the methylome of Alphaproteobacteria.

Our initial validation with *Caulobacter* showed the characteristic cell cycle dependent methylation of GANTC sites due to restricted activity of the CcrM methyltransferase only to the end of the replication cycle. The correlation with previous measurement with SMRT sequencing approaches was very high (SFig 1), supporting this approach. We then tested the assumption that loss of the Lon protease results in dysregulation of CcrM activity because Lon normally degrades CcrM. Consistent with this model, Δ*lon* strains fail to show the same cell cycle regulation of GANTC methylation and their genomes are generally more methylated. Next, we examined if AlkB, a dealkylase capable of removing m6A marks from DNA *in vitro* (^13^), affected methylation *in vivo*. We found no evidence for a global role of AlkB in DNA methylation *in vivo* under normal growth conditions, but identified some sites that were differentially methylated in an Δ*alkB* strain that could impact gene expression, as shown with CcrM expression (Figure 2).

For *Caulobacter* Δ*lon*, we found a handful of sites with particularly substantial changes in methylation and were also altered in transcription in this strain. The gene CCNA_00169 has multiple GANTC sites in its promoter region all of which were hypermethylated in Δ*lon* and showed an increased transcription in Δ*lon*. Although this gene is annotated as a hypothetical lipoprotein, it is adjacent to a TonB receptor and next to a cluster of Type 2 secretion genes, suggesting that CCNA_00169 may be a Type 2 secreted substrate (^29^). Loss of this gene has a general fitness defect across multiple conditions based on bar-seq results and is highly conserved (^30^). Interestingly, this same region is normally kept hypomethylated due to protection by MucR ( ^11^), supporting a possible link between Lon and MucR in *Caulobacter* either directly or due to stabilization of CcrM.

Using this nanopore approach, we measured the methylome of *Brucella abortus* for the first time that we know of. As would be predicted from the conservation of CcrM, we find that GANTC methylation shows the same cell cycle dependency as *Caulobacter.* This dependency is found in both chromosomes of *Brucella*. Interestingly, loss of Lon protease in *Brucella* has species-specific effects on CcrM-dependent methylation. Cell cycle dependency of methylation was lost in Chromosome I in Δ*lon*, much like that seen in *Caulobacter*; however, Chromosome II showed the same methylation pattern as in wild-type. Additionally global GANTC methylation was lower in Δ*lon* than in wild-type for both chromosomes, in contrast to our expected findings in *Caulobacter* (Figure 3), leading to the possibility that steady state levels of CcrM may be overall lower in a Δ*lon* strain. We speculate that CcrM cell cycle regulation must be altered for Chromosome I to be affected, consistent with a role of Lon in regulating CcrM (^8,21^). The Lon-independence of this methylation in Chromosome II suggests an additional layer of control where another methyltransferase may be present to enforce this mark or that the pool of CcrM associated with Chromosome II is still degraded even in the absence of Lon. Given that Lon is important for *Brucella* virulence (^16,31^) this complex connection with CcrM methylation will be a future topic of interest.

The transcription factor MucR binds to *Brucella’s* chromosomes in an H-NS like manner, is recruited to AT-rich regions, and may contribute to nucleoid structure (^18^). Our findings suggest that MucR also protects specific hypomethylated sites from methylation by CcrM. One of these sites on Chromosome II is adjacent to BAB2_1107, a known autotransporter important for adhesion to host cells (^32^). Of note, is that loss of MucR affects Chromosome II only in short-range interactions, versus its more global effect on Chromosome I (^18^). Similarly, loss of Lon affects Chromosome I methylation more dramatically (Figure 3) suggesting a possible link between MucR and Lon. Finally, an equal number of hypomethylated sites remain protected in Δ*mucR* and are not bound by MucR (Figure 4), suggesting that other MucR-like proteins may shield those sites. However, overall, loss of MucR does not dramatically affect the global methylation patterns of either of the chromosomes.

Thanks to new machine learning pipelines such as mCaller (^20^) or Dorado (^25^), together with the ever decreasing costs of nanopore sequencing, detection of epigenetic modifications can become more standardized. In our work presented here, we focused on two Alphaproteobacteria genomes and monitor how known or predicted regulators of methylation affect their epigenome. However, we note that the increasing adoption of nanopore sequencing of plasmids by contract companies (Plasmidsaurus, etc.) will make monitoring epigenetic changes in these genetic elements an unexpected bonus.

## Methods

### Bacterial growth conditions and sequencing

Refer to Supplementary Table 1 for all strains used in this paper. *Caulobacter crescentus* NA1000, along with its derivative strains, were initially cultured on Peptone Yeast Extract (PYE) agar plates. These plates contain a specific nutrient mix: 2 g/L of peptone, 1 g/L of yeast extract, 0.5 mM of MgSO4, 0.5 mM of CaCl2, and 1.5% agar (produced by Difco), and were incubated overnight at 30°C. Following incubation, isolated colonies were picked and further cultured in PYE liquid media overnight at 30°C within shaken flasks. The cultures were then diluted to achieve a starting optical density (OD600) of 0.1, and allowed to grow to a mid-exponential phase (achieving an OD600 of 0.6) prior to the isolation of chromosomal DNA.

*Brucella abortus* 2308 and derivative strains were grown on Schaedler agar (Becton, Dickinson, and Co., Sparks, MD, USA) supplemented with 5% defibrinated bovine blood (SBA) incubated at 37°C under 5% CO2 or in Brucella Broth (Becton, Dickinson, and Co., Sparks, MD, USA) incubated at 37°C with shaking. Growth media were supplemented with kanamycin (45 ug/mL) or chloramphenicol (10 ug/mL) when appropriate. For isolation of chromosomal DNA, *Brucella abortus* 2308 and derivative strains were grown overnight in Brucella Broth, then subcultured to an OD 600 of 0.1 and then grown to mid-exponential phase (OD600 of 0.6) before isolation of chromosomal DNA by guanidine thiocyanate as described previously (^33^). Genomic DNA was sequenced by Seqcenter, LLC. *Caulobacter* strains were sequenced using nanopore R.9.5 pores, and *Brucella* strains were sequenced using nanopore R.10.4.1 pores.

### Western Blot Analysis

For western blot analysis, *Caulobacter* strains were grown overnight in PYE media, and back-diluted to an OD600 of 0.05. Then grown in shaking flasks at 30°C in PYE, samples were collected at 1.5, 3, 6, and 9 hours and centrifuged at 26,000 rpm for 10 minutes to pellet the cells. The pellets were resuspended in 2x SDS-PAGE dye (12% glycerol, 4% SDS, 100 mM Tris-HCl pH 6.8, trace bromophenol blue, 40 mM DTT). 10µl of each sample was loaded into a 16-well 10% Bis-Tris plate for electrophoresis. Post-separation the membrane was incubated overnight with CcrM antibodies at a 1:5,000 dilution in 5% milk. Following overnight primary probing at 4 °C, the membranes were washed three times with Tris Buffered Saline with 1% Tween (TBST). Proteins were then visualized using IRdye-labeled mouse anti-rabbit antibody (LI-COR Biosciences) at 1:10,000 dilution and an Odyssey Scanning system (LI-COR). As ClpP levels remain constant in *Caulobacter*, it was used as a loading control for quantification.

Membranes were incubated for 1 hour with anti-ClpP (1:5000 dilution) and visualized using IRdye-labeled mouse anti-rabbit antibody (LI-COR Biosciences) at 1:10,000 dilution and an Odyssey Scanning system (LI-COR).

### Pipeline for nanopore R.9.5 through mCaller

Raw nanopore sequencing outputs (Fast5 format) were quality-controlled and trimmed using Guppy (https://community.nanoporetech.com). High-quality reads were aligned to the reference genome NC_011916.1 (^34^) with Minimap2 (^35,36^), using default parameters. Fast5 files were converted to slow5 files (^37,38^) and aligned to the mapped reads using the ‘eventalign’ function within Nanopolish (^39^). Methylated bases were quantified using mCaller (^20^), employing the precompiled ‘r95_twobase_model_NN_6_m6A.pkl’ model for assessing modified adenines. The methylation results were then annotated using Bedtools (^40^) and then analyzed with R (^41^).

### Pipeline for nanopore R.10.4.1 through Dorado

Raw nanopore sequencing outputs (Fast5 format) from nanopore sequencing, were converted into pod5 files for optimization. Dorado’s (^25^) standard basecalling command was employed, utilizing the model ‘dna_r10.4.1_e8.2_400bps_sup@ v4.2.0’. In addition, modified base calling was executed using the specialized model ‘dna_r10.4.1_e8.2_400bps_sup @v4.2.0_6mA@v2’. The produced bed files were then aligned to NCBI reference genomes NC_007618.1 and NC_007624.1 (^42^) with the mapped reads using the align and sort functions of Samtools (^43^). The resulting modified bam files were run with modbam2bed (^44^) with default settings and then analyzed with R (^41^).

### Data Processing in R

#### Definitions in relation to methylation

In our study, motifs are defined as individual occurrences of the ‘GANTC’ sequence. Given the palindromic nature of ‘GANTC’, we pair motifs within a 2-base-pair distance to define “sites”.

When comparing the probability of methylation among different strains, we considered motifs that deviated greater than or less than 2.5 standard deviations from the mean of a dataset as significantly different. Expanding upon this, sites in which both motifs fell below 2.5 standard deviations below the mean methylation probability for that particular strain were classified as hypomethylated.

#### Identification of and filtration of m6A motifs

Data was imported from both mCaller and Dorado pipelines into RStudios (^45^) for analysis. Probability of methylation of GANTC motifs obtained through obtaining positional index through regexes of reference files and filtration of bed files. Additional motifs were identified through filtering all adenines with above a 80% score, coverage greater than 50, and homology with previously identified methyltransferase motifs in the Rebase database. All identified motifs in this study are available in Supplementary Table 3.

Slopes of probability of methylation as function of genomic position for each strain’s probability of methylation was calculated by finding the linear regression between genomic position from both ends of the origin to the terminus under the assumption that there is symmetry over the terminus. Pearson correlation coefficient between Nanopore and SMRT sequencing data was performed using data obtained from (^10^), full methods described with Supplementary Figure 1. Additional methods, and scripts are available on GitHub (https://github.com/mbcampbell-work/Alphaproteobacteria-Methylation).

#### Identification of genes in relation to gene expression

In order to assess whether changes in methylation had association with transcriptional changes, we conducted Fisher’s exact tests comparing the change of GANTC methylation upstream of genes against transcriptional change in mutants. Focusing upon GANTC sites located in intergenic regions, particularly those within 300 base pairs of a gene’s start codon or within 200 base pairs of a known transcription start site (TSS). TSS data for *Caulobacter* were retrieved from prior study available at (^46^).

Changes in gene expression between the Δ*lon* mutant and wild-type *Caulobacter* were analyzed using data from the Gene Expression Omnibus (GEO) database, under accession number GSE152426, as reported in (^23^). Additionally, microarray and ChIP-seq data in relation to the Δ*mucR* mutant of *Brucella abortus* were sourced from (^18^).

## Acknowledgements

We gratefully acknowledge Tommy Tashjian, who initiated the methylation project in *Caulobacter* and generated the Δ*alkB* strain. We also thank all members of the Chien lab for comments and discussion. Funding for this work was provided by NIH (NIGMS R35GM130320 (PC), NIAID R21AI148132 (MR), NIAID R01AI172822 (MR)), and USDA-NIFA 7002581 (PC).

## Supplementary Material

### Supplementary Text and Figures

**SFigure 1:** Correlation between Nanopore and SMRT sequencing

**SFigure 2:** Distribution of GANTC sites among Caulobacterales and Rhizobiales

**SFigure 3:** Methylomes of *Brucella* strains in stationary phase

**SFigure 4:** Methylomes of non-GANTC motifs in *Caulobacter*

**SFigure 5:** Methylomes of non-GANTC motifs in *Brucella*

### Supplementary Tables

**STable 1:** Strains used in this study

**Stable 2:** Distribution of GANTC motifs among Alphaproteobacteria with CcrM homologs

**STable 3:** m6A motifs identified in *Caulobacter crescentus* and *Brucella abortus*

**STable 4:** GANTC sites that show both significant changing in methylation and transcriptional change between Δ*lon* and wild-type *Caulobacter crescentus*

**STable 5:** GANTC sites that show both significant changing in methylation between Δ*alkB* and wild-type *Caulobacter crescentus*

**STable 6:** Hypomethylated sites in wild-type *Brucella abortus*

### Supplementary Files

**SFile 1:** Exponential *Caulobacter crescentus* NA1000 methylation calls (mCaller)

**SFile 2:** Exponential *Brucella abortus* 2308 methylation calls (dorado)

**SFile 3:** Stationary phase *Brucella abortus* 2308 methylation calls (mCaller)

### GEO

Additional data is available at GEO xxx.

### SRA

Raw FAST5 files of all strains are available at SRA xxx.

### GitHub

All computational methods and scripts are available at https://github.com/mbcampbell-work/Alphaproteobacteria-Methylation.

### Supplementary Text

#### Comparison of Nanopore and SMRT sequencing in methylation detection

A pivotal aspect of our study revolves around the methodologies employed for m6A methylation detection. Although there is an the absence of a gold standard for m6A methylation, akin to bisulfite conversion for 5mC methylation ( ^48^), current technology includes developing platforms based nanopore sequencing and SMRT sequencing (^49^).

Single Molecule, Real-Time (SMRT) sequencing, developed by Pacific Biosciences, detects methylation by analyzing DNA polymerase kinetics during the sequencing process ( ^50,51^). As the polymerase incorporates nucleotides into the DNA strand being synthesized, the presence of methylated bases causes distinctive changes in the enzyme’s speed, which are recorded as variations in the inter-pulse duration (IDP) of fluorescent signals (^50^).

In contrast, Nanopore sequencing, developed by Oxford Nanopore Technologies, detects methylation and other DNA modifications directly during the sequencing process (^52^). This technique involves threading single DNA molecules through a nanopore embedded in a synthetic membrane. As the DNA passes through the pore, changes in the ionic current are measured, with different bases and modifications (such as methylation) causing distinct patterns of disruption in the current.

Our choice of utilizing nanopore sequencing allows us to compare the results we saw with prior work which has utilized SMRT sequencing for methylation analysis in wild type *Caulobacter* (^10^), in which they were able to calculate methylation by sectioning the genome off into 20 kb sections and calculating the average IPDratio per section. By implementing a similar process of averaging probability of methylation across a range of 1 to 200kb sections of genome, we found a strong correlation between the two metrics: IPDratio (SMRT sequencing) and Probability of Methylation (Nanopore sequencing) (SFig 1). However, it was of note that comparing individual sites showed a substantially weaker correlation.

This observation suggests two possible interpretations: it may reflect inherent variability in methylation patterns, or it may indicate that the methodological differences in how each technique identifies methylation lead to variations in precision at a localized scale. However, by aggregating methylation calls over larger genomic regions, we were able to maintain consistent detection of methylation patterns across both nanopore and SMRT sequencing approaches. This consistency suggests that, despite their differences, both methods are reliable for identifying broad methylation patterns.

#### GANTC distribution

Previous studies have systematically studied the conservation of CcrM homologs among Alphaproteobacteria, as well as the distribution and placement of GANTC motifs (^10^).

Highlighting that GANTC motifs are less frequent in the genome than expected compared to a random sequence of penta-nucleotides. However there appears to be an underlying selection that favors the placement of GANTC within intergenic regions that highlights the role these sites may play in the epigenetic regulation of genes. Using BLAST (^53^) to identify known CcrM homologs across the orders Caulobacterales and Rhizobiales, and obtaining reference genomes from the NCBI database (^54^) we were able to calculate the ratio of counted to expected number of GANTC’s for a selected region (script available on linked GitHub). Our analysis aligns with prior findings (SFig 2). The differences in number of GANTC sites found between *Brucella abortus* 2308 and *Caulobacter crescentus NA1000* is likely a result of differences in GC content of the respective genomes.

#### Stationary phase Brucella

The relationship between *Brucella* virulence and its growth in the stationary phase has been well characterized ( ^55^). *Brucella* exhibits a distinct stationary-like growth pattern post-infection in murine hosts, where it reaches a plateau of growth and shows minimal net replication ( ^26^).

Given that CcrM is cell cycle dependent, and stationary phase is defined by its limited replication we expected to see a significant change in our stationary phase methylomes (SFig 3) compared to our exponential phase methylome (Figure 3. A-C). In alignment with expectation, we found that all strains of *Brucella* in stationary exhibited a narrow range of change in methylation across Chromosome I, and flattened slope for Chromosome 2.

#### Non-GANTC motifs

Beyond the recognized m6A motif, GANTC, two additional m6A motifs have previously been identified in *Caulobacter*, 5’-CGACC**<u>A</u>**G-3’ and 5′-CG**<u>A</u>**C(N7)TRGG-3′, which have been linked to restriction modification systems (^9^). Analysis of non-GANTC m6A methylation probabilities across the genome (SFig 4) indicates an absence of cell cycle-dependent variations, reinforcing the idea that CcrM operates as the sole cell cycle-regulated adenine methyltransferase.

Moreover, when analyzing non-GANTC m6A marks in Δ*alkB*, no significant differences emerge, further suggesting that AlkB does not significantly influence the removal of methyl groups from m6A methylation in vivo. While the motif 5′-CG**<u>A</u>**C(N7)TRGG-3′ consistently exhibits high levels of methylation, aligning with previous observations (^9^), the low methylation probability associated with 5’-CGACC**<u>A</u>**G-3’ is likely not indicative of a biological effect but rather reflects mCaller’s reduced confidence in motifs that significantly deviate from its pre-trained model.

In our analysis of adenines not associated with GANTC motifs on Chromosome I of *Brucella*, we identified 260 instances of potential m6A sites that met our criteria: a score above 80%, coverage greater than 50 reads, and probability of methylation greater than 0.33 (Our definition of hypomethylated in wild-type *Brucella*). Out of these 260 instances, 209 were linked to the m6A motif 5′-**<u>A</u>**CCNNNNNGTCG-3′. This motif, as inferred from homology with previously identified methyltransferases in the Rebase database, is associated with the R-M methyltransferase HsdM. A similar pattern emerged on Chromosome II of *Brucella*, where 108 out of 120 adenines which met our criteria were also related to the HsdM motif. Across all studied strains, we observed neither cell cycle dependence nor any significant changes in methylation patterns (SFig 5).

## Supplementary Figures

**Supplementary Figure 1:**
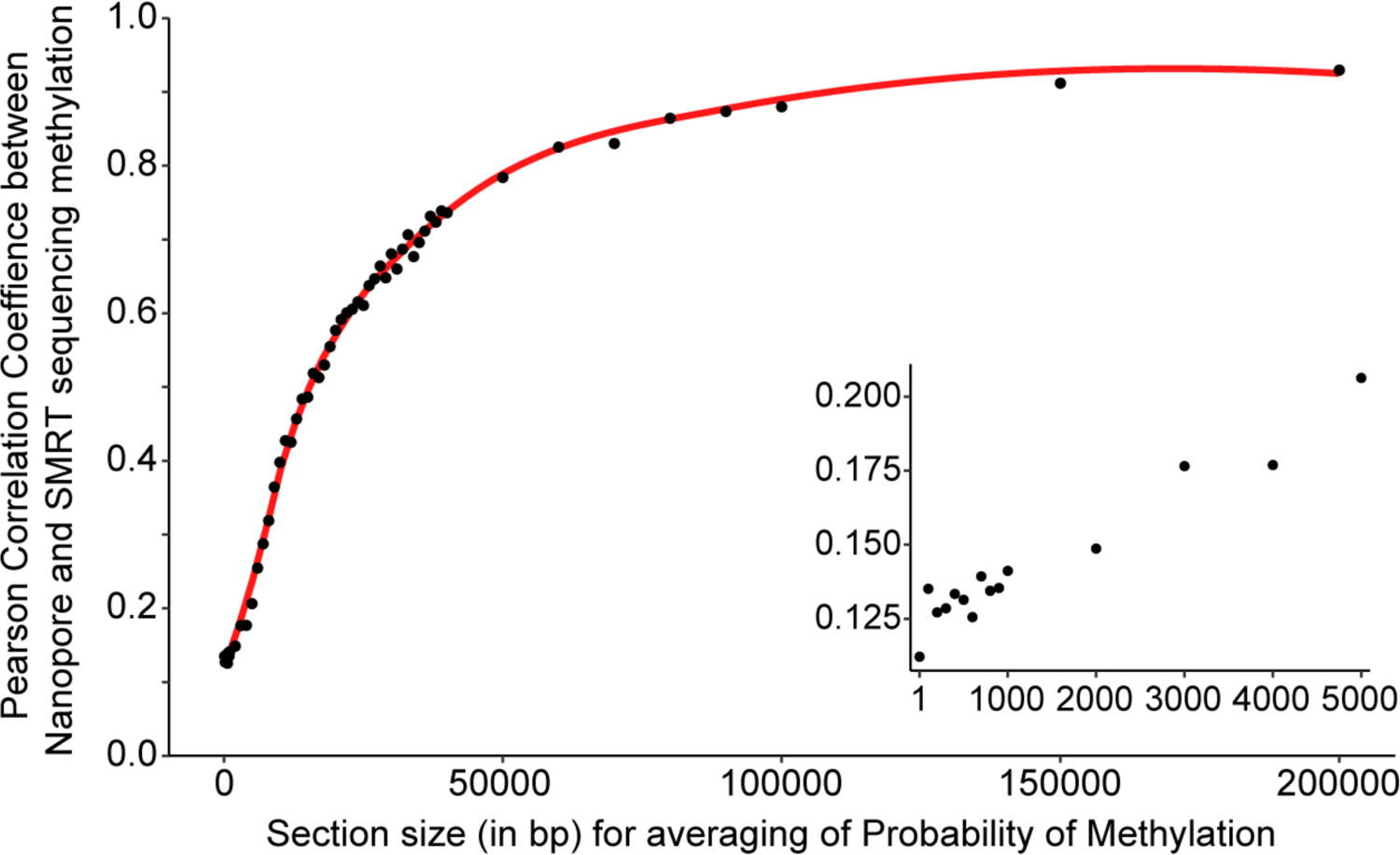
Comparison of Nanopore and SMRT sequencing in methylation detection Correlation between Nanopore and SMRT sequencing metrics for methylation detection in wild-type *Caulobacter* as a process of averaging calls over x (between 1 and 200,0000) base-pairs. Pearson correlation coefficients ranged from 0.11 to 0.93.

**Supplementary Figure 2:**
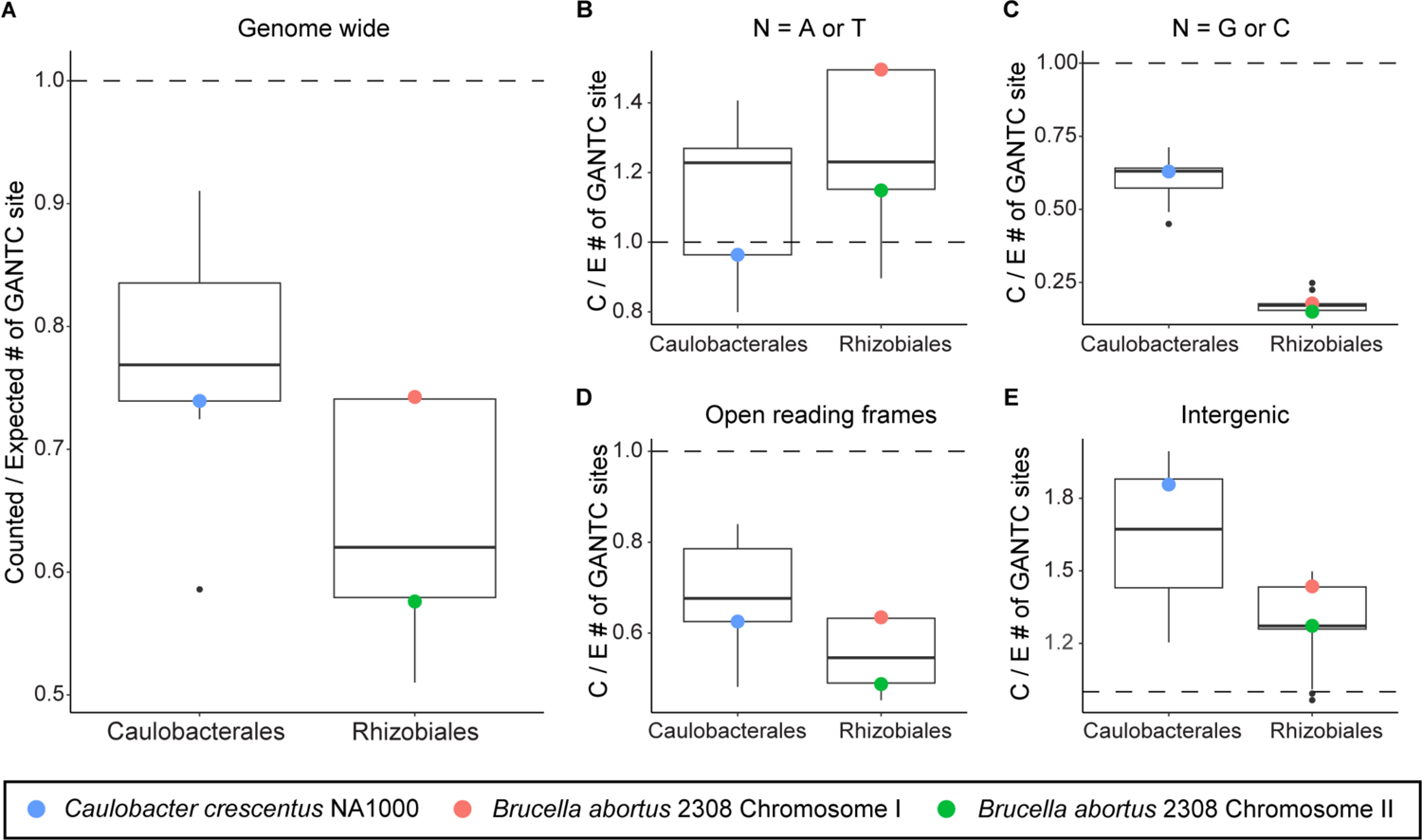
(**A-E**) Boxplots of the ratio of counted to expected (C/E) GANTC motifs across various genomic contexts for Caulobacterales and Rhizobiales: genome-wide (**A**), when the ambiguous base (N) in the GANTC sequence represents adenine or thymine (**B**), and cytosine or guanine (**C**), in open reading frames (**D**), and intergenic regions (**E**). The dataset encompasses *Caulobacter crescentus* NA1000 (highlighted in blue) and nine other Caulobacterales with identified CcrM homologs via BLAST. As well as *Brucella abortus* 2308 (Chromosome I in green and Chromosome II in red) and thirteen additional Rhizobiales also identified to contain CcrM homologs through BLAST.

**Supplementary Figure 3:**
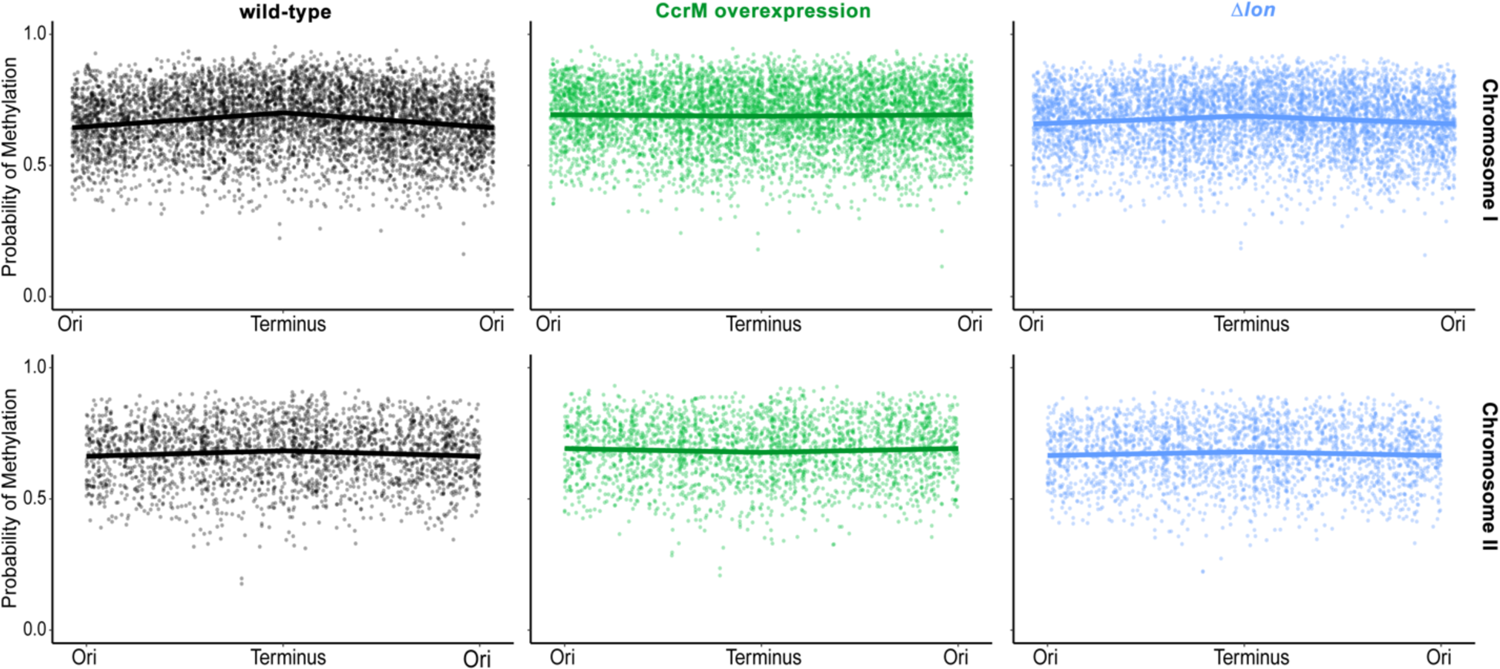
Methylomes of *Brucella* strains in stationary phase Probability of CcrM-dependent methylation at GANTC motifs on Chromosome I (upper graphs) and Chromosome II (lower graphs) of stationary phase *Brucella* in different genetic contexts: wild-type (black), with CcrM overexpression (green), and deletion of *lon* (blue).

**Supplementary Figure 4:**
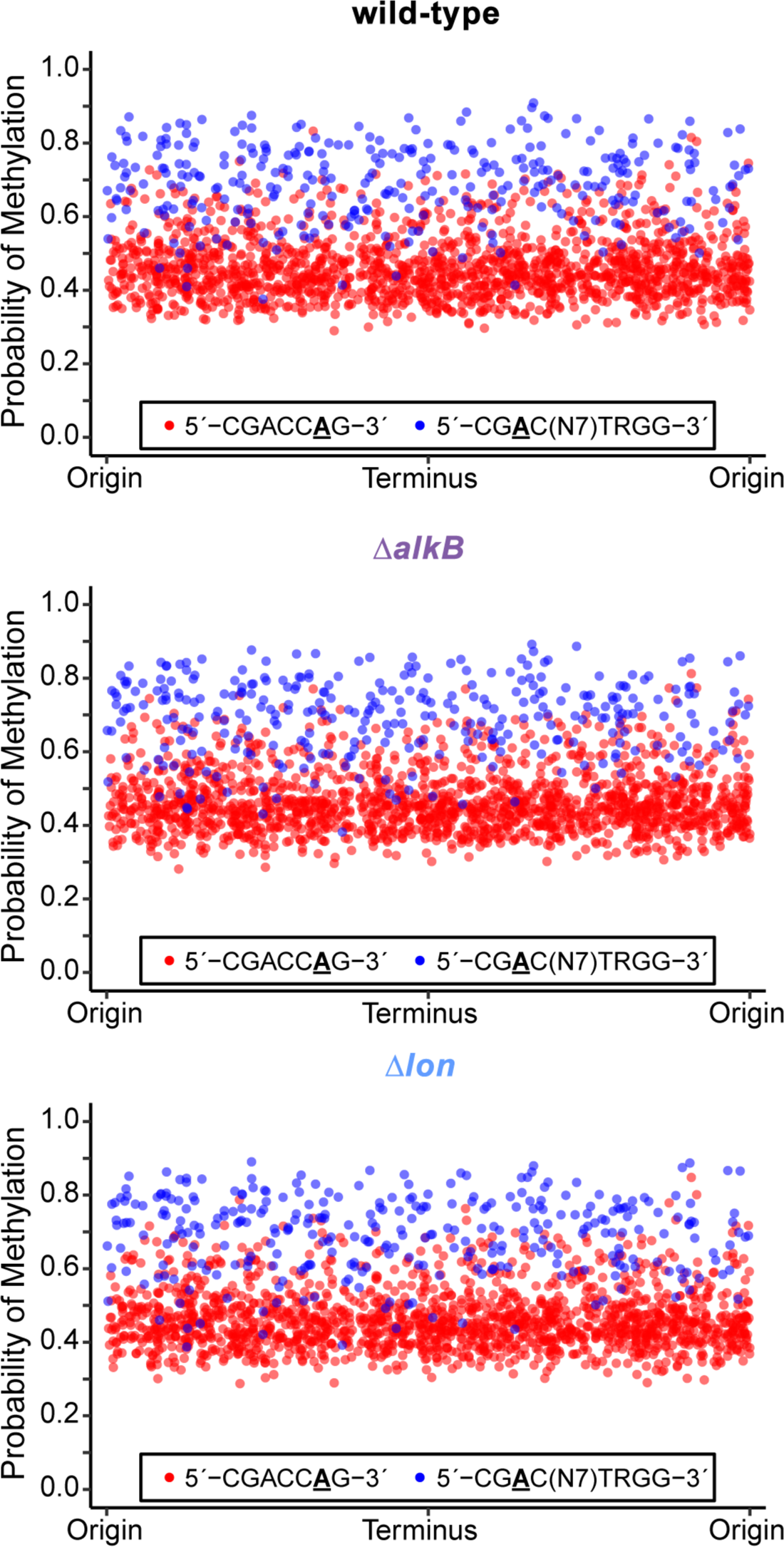
Methylomes of non-GANTC m6A motifs in *Caulobacter* strains. Probability of non-CcrM-dependent m6A methylation at both 5’-CGACCAG-3’ (red) and 5′-CGAC(N7)TRGG-3′ (blue) motifs on *Caulobacter* in different genetic contexts: wild-type (top), deletion of *alkB* (middle), and deletion of *lon* (bottom).

**Supplementary Figure 5:**
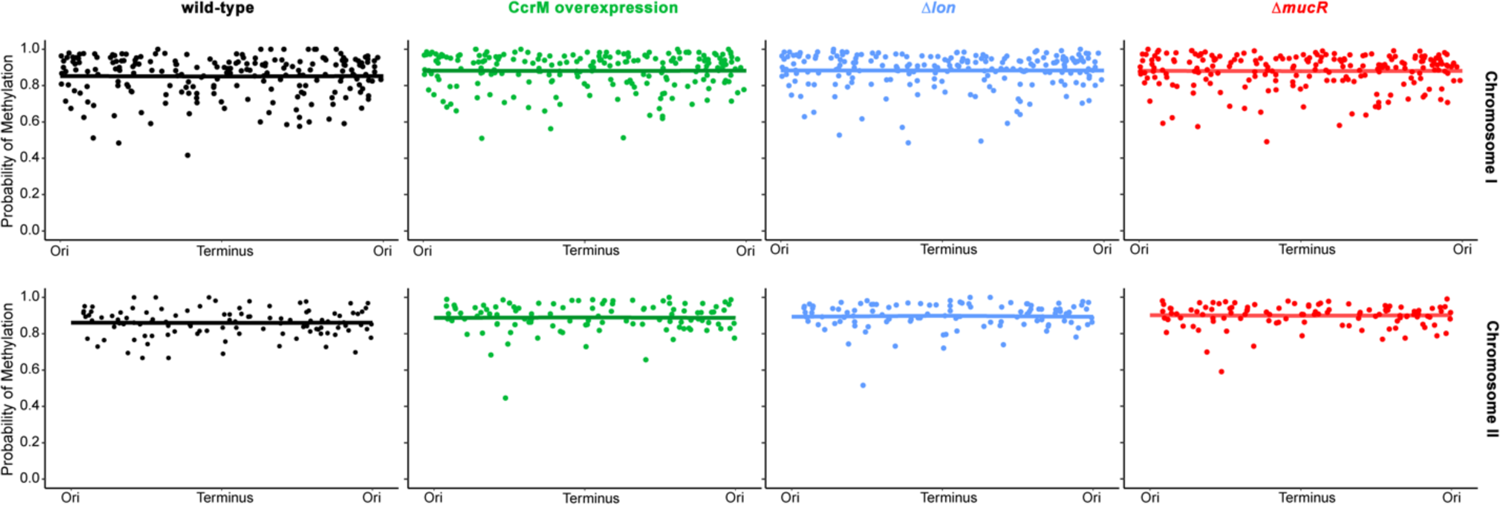
Methylomes of non-GANTC motifs in *Brucella* strains Probability of HsdM-dependent methylation at 5′-ACCNNNNNGTCG-3’ motifs on Chromosome I (upper graphs) and Chromosome II (lower graphs) of *Brucella abortus* in different genetic contexts: wild-type (black), with CcrM overexpression (green), deletion of *lon* (blue), and deletion of *mucR* (red).

